# The autophagy receptor Ncoa4 controls PPARγ activity and thermogenesis in brown adipose tissue

**DOI:** 10.1101/2025.02.02.636110

**Authors:** Leslie A. Rowland, Kaltinaitis B. Santos, Adilson Guilherme, Sean Munroe, Lawrence M. Lifshitz, Sarah Nicoloro, Hui Wang, Matthew F. Yee, Michael P. Czech

## Abstract

Adipose tissue dysfunction leads to a variety of deleterious systemic consequences including ectopic lipid deposition and impaired insulin sensitivity. PPARγ is a major regulator of adipocyte differentiation and functionality and is thus a determinant of systemic metabolic health. We recently reported that deletion of adipocyte fatty acid synthase (AdFasnKO) impairs autophagy in association with a striking upregulation of genes controlled by PPARγ, including thermogenic uncoupling protein 1 (Ucp1). In this present study, screening for PPARγ coactivators regulated by autophagy revealed a protein denoted as Nuclear receptor coactivator 4 (Ncoa4), known to mediate ferritinophagy and interact with PPARγ and other nuclear receptors. Indeed, we found Ncoa4 is upregulated in the early phase of adipocyte differentiation and is required for adipogenesis. Ncoa4 is also elevated in FasnKO adipocytes and necessary for full upregulation of Ucp1 expression *in vitro*, even in response to norepinephrine. Consistent with these findings, adipose-selective knockout of Ncoa4 (AdNcoa4KO mice) impairs Ucp1 expression in brown adipose tissue and cold-induced thermogenesis. Adipose-selective double KO of Fasn plus Ncoa4 (AdFasnNcoa4DKO mice) prevents the upregulation of classic PPARγ target genes normally observed in the white adipose tissue of AdFasnKO mice, but not thermogenic Ucp1 expression. These findings reveal Ncoa4 is a novel determinant of adipocyte PPARγ activity and regulator of white and brown adipocyte biology and suggest that manipulation of autophagy flux modulates PPARγ activity and key adipocyte functions via Ncoa4 actions.

## Introduction

Adipose tissue is a major contributor to metabolic health, due in part to the ability of adipocytes to safely store and sequester lipids, protecting other organs from lipotoxicity. In addition, beige and brown adipose tissues can improve metabolism by increasing energy expenditure and fuel utilization, and all adipose tissues secrete beneficial factors that target other organs to promote metabolic health [1,2]. A major transcription factor responsible for controlling many of these adipocyte functions is PPARγ, which directs gene programs involved in adipogenesis, lipid storage, glucose uptake, and thermogenesis [3]. Accordingly, manipulation of PPARγ activity in adipocytes can strongly affect whole-body metabolism, thermogenesis, and insulin sensitivity [4–7].

PPARγ activity is highly regulated and can be modulated through posttranslational modifications, coregulator binding, and degradation of PPARγ itself via the proteasome [6,8–10]. More recently, it’s been shown that PPARγ activity can also be regulated by autophagy. Coactivators of PPARγ, including Ncoa1/SRC1 and Ncoa2/TIF2 are degraded in an autophagy-dependent manner, and their loss is associated with decreased PPARγ action, resulting in reduced adipogenic gene expression and dysfunctional adipocytes [11,12]. In brown adipocytes, the autophagic degradation of the PPARγ corepressor Ncor1 is required to maintain PPARγ activity and brown adipocyte functionality [13]. Thus, the extent of autophagic flux can determine adipocyte PPARγ activity via coregulator turnover and appears to occur in a context and tissue-specific manner.

In addition to controlling PPARγ activity, autophagy has been shown to affect other adipocyte functions, including the development of beige adipose tissue. Autophagy impairment in adipocytes using the ap2-Cre-driven Atg7 knockout mouse model led to the development of beige-like multilocular white adipocytes, increased brown fat mass, and improved whole-body insulin sensitivity [14,15]. Deletion of another autophagy regulator, Atg5, in adipocytes prevented the whitening, or reversal, of beiging, indicating beige adipocyte maintenance is also controlled by autophagy [16]. Since PPARγ directly controls thermogenic gene expression in white and brown adipocytes, it’s possible some of these beiging effects could be due to increased PPARγ activity as a result of altered autophagic turnover of PPARγ coregulators, though this has not been explored.

In recent years, the adipocyte-specific fatty acid synthase (Fasn) knockout (AdFasnKO) mouse model has been studied to reveal new mechanisms of adipocyte beiging. These mice lack the ability to synthesize fatty acids *de novo* in adipocytes due to the knockout of this last enzyme in the *de novo* lipogenesis pathway. Unexpectedly, these mice exhibit strong beiging of the white adipose tissue and improved glucose homeostasis [17–19]. We found that adipose tissue macrophages are associated with this beiging; however, the changes *within* the AdFasnKO adipocytes that led to beiging remained unknown [20]. Recently, we found that AdFasnKO adipocytes have a significant impairment in autophagy, due to defects in autophagosome-lysosome fusion and lysosomal dysfunction [21]. Due to the known roles of autophagy in PPARγ regulation and beige adipose tissue formation, we hypothesized that the dysfunctional autophagy in AdFasnKO adipocytes resulted in the beiging phenotype.

In this present study, using the adipocyte-specific FasnKO mouse model, we identify a previously unknown role for the autophagy receptor Ncoa4 in adipocyte biology. We find that Ncoa4 is necessary to drive PPARγ-directed differentiation of progenitor cells to adipocytes in culture and for driving PPARγ dependent genes in white adipose tissue. Importantly, this novel cofactor Ncoa4 is required for optimal thermogenesis in brown adipocytes *in vivo,* as demonstrated in mice with adipose-selective Ncoa4 depletion.

## Results

### PPARγ target gene expression is increased in AdFasnKO adipocytes

To understand gene expression changes in the AdFasnKO adipocytes, which may lead to beiging of the previously described AdFasnKO mouse white adipose tissue (WAT), we analyzed bulk RNA sequencing data from the subcutaneous WAT of these mice [19]. Specifically, we used the TRUSST database to identify transcription factors predicted to contribute to the gene expression changes [22]. Interestingly, PPARγ was the most significant transcription factor predicted to drive the genes upregulated by AdFasnKO (Figure 1a). Strikingly, using the PPARgene database [23], we found that of the 35 experimentally verified PPARγ target genes expressed in mouse adipose tissue and in our RNAseq dataset from AdFasnKO mice, 29 (83%) were significantly upregulated with padj < 0.05 (or 32 (91%) with padj <0.1) (Figure 1b). It is known that PPARγ regulates thermogenic gene expression, which is also controlled by other pathways, namely the cAMP/PKA pathway. To ensure the enrichment in PPARγ target genes in the AdFasnKO adipose tissue was not only a reflection of the increased thermogenic gene expression due to the activation of cAMP/PKA pathway or others, we confirmed the upregulation of the PPARγ target gene Fabp4 in the gonadal white adipose tissue of AdFasnKO male mice, a tissue that does not exhibit increased thermogenic gene expression (Figure 1c and 1d).

**Figure 1.**
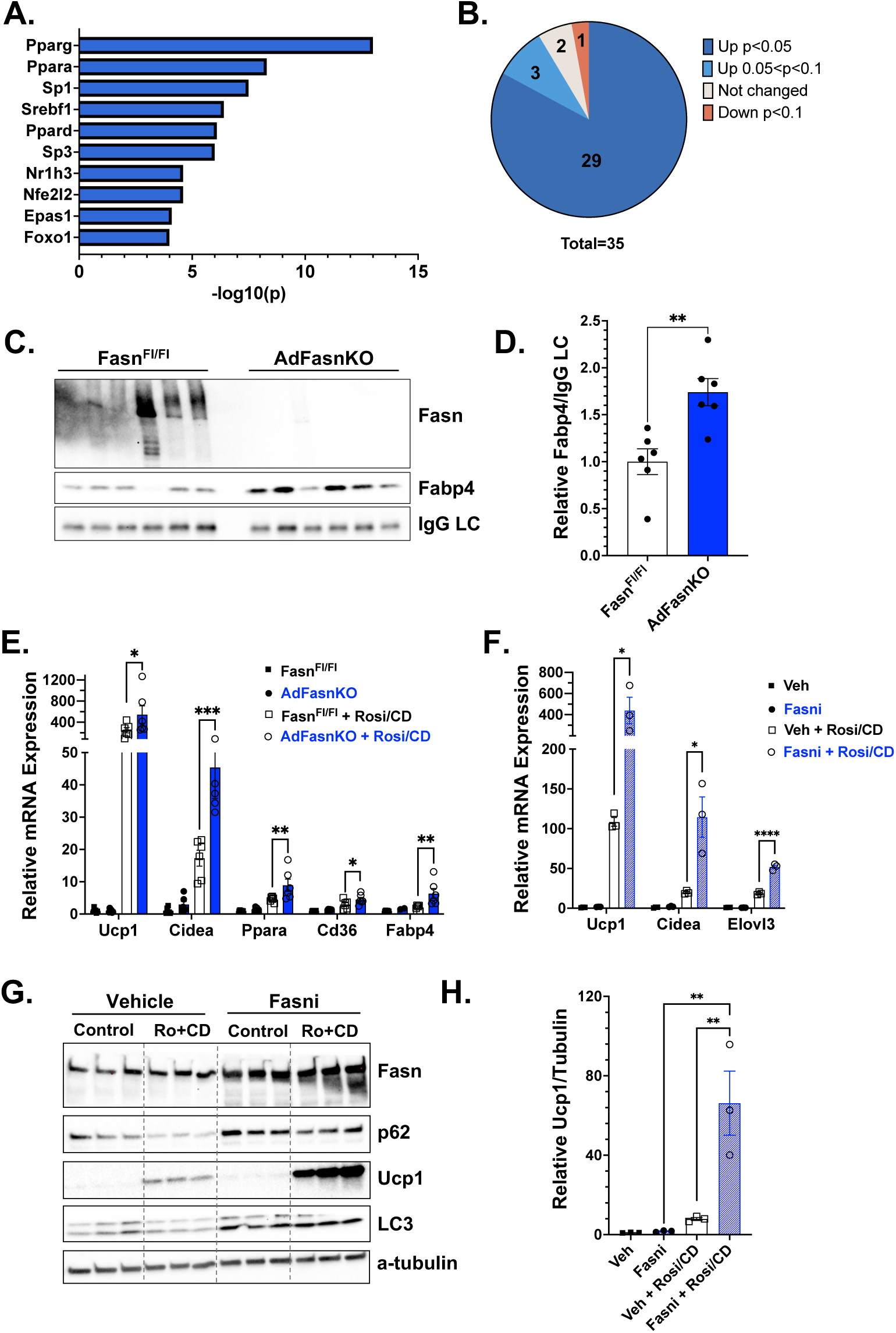
Loss of Fasn in adipocytes potentiates PPARγ-driven gene expression. A) TRUSST analysis of genes upregulated (padj<0.05, FC>1) in AdFasnKO subcutaneous white adipose tissue (WAT). From RNA-sequencing analysis in Guilherme et al. 2023 [19]. B) Numbers of PPARγ target genes, as annotated by the PPARgene database, upregulated (Up), not changed, or downregulated (Down) in AdFasnKO WAT. C) Western blot for Fasn, Fabp4, and mouse IgG LC (light chain) in gonadal WAT of AdFasnKO mice. D) Quantification of Fabp4 in western blot in C. Student’s t test ** p<0.01 E) Quantitative PCR (qPCR) analysis of PPARγ-regulated genes in *in vitro* differentiated Fasn^Fl/Fl^ and AdFasnKO adipocytes treated with 50nM rosiglitazone (Rosi) and 50nM CD3254 (CD) or DMSO for 72 hrs. 2-way ANOVA with Sidak’s multiple comparisons test * p<0.05, ** p<0.01, *** p<0.001 F) qPCR analysis of PPARγ-regulated thermogenic genes in wild-type *in vitro* differentiated adipocytes treated with 50nM rosiglitazone (Rosi) and 50nM CD3254 (CD) or DMSO with or without 100nM TVB-3664 (Fasni) for 48 hrs. 2-way ANOVA with Sidak’s multiple comparisons test * p<0.05, **** p <0.0001 G) Western blot of adipocytes treated as in F. H) Quantification of Ucp1 in the western blot in G. 2-way ANOVA with Sidak’s multiple comparisons test ** p<0.01.

### Loss of Fasn activity potentiates PPARγ-driven gene expression

To confirm the changes in PPARγ gene expression occurred in adipocytes, rather than other cell types present in the adipose tissue, we isolated preadipocytes from AdFasnKO subcutaneous adipose tissue and initiated their differentiation *in vitro*. In unstimulated conditions, AdFasnKO adipocytes did not differ from controls in PPARγ target gene expression. We then stimulated the cells with the PPARγ agonist rosiglitazone, in combination with the RXRα agonist CD3254 since PPARγ functions as an obligate heterodimer with RXRα. Under this latter condition, AdFasnKO adipocytes showed significantly higher PPARγ-dependent gene expression (Figure 1e). Treatment of fully differentiated wild-type adipocytes with a Fasn inhibitor (Fasni) similarly potentiated PPARγ-dependent gene expression in the presence of PPARγ:RXRα agonists (Figure 1f). Furthermore, these changes were evident at the protein level, as Ucp1 protein was significantly elevated in the presence of the Fasn inhibitor (Figure 1g and 1h). Together, these data indicate that a loss of Fasn activity in adipocytes potentiates ligand-dependent, PPARγ:RXRα-driven thermogenic and adipogenic gene expression.

### Known autophagy-regulated PPARγ coactivators Ncoa1 and Ncoa2 are not changed in FasnKO adipocytes

The studies described above indicated that the upregulation of PPARγ target gene expression was dependent on the presence of ligands for PPARγ:RXRα in cultured adipocytes. This suggested that a PPARγ coactivator may be upregulated in the FasnKO adipocytes, since nuclear receptor regulators (coactivators) often bind or activate transcription in a ligand-dependent manner. It’s recently been shown that levels of the PPARγ coactivators Ncoa1/SRC1 and Ncoa2/TIF2 are controlled by autophagy [11,12], thus we reasoned they may be upregulated in the autophagy-impaired FasnKO adipocytes. However, neither Ncoa1/SRC1 nor Ncoa2/TIF2 were affected in AdFasnKO adipose tissue in either fed or fasting conditions, though autophagy was impaired as indicated by increased p62 (Figure 2a and 2b). Moreover, in differentiated adipocytes, Fasn inhibition did not significantly affect Ncoa1/SRC1 or Ncoa2/TIF2 levels, despite autophagy inhibition (increased p62) (Figure 2c and 2d). We also found that expression of PRDM16, a PPARγ coactivator important for thermogenic gene expression [24], was not changed in FasnKO adipose tissue.

**Figure 2.**
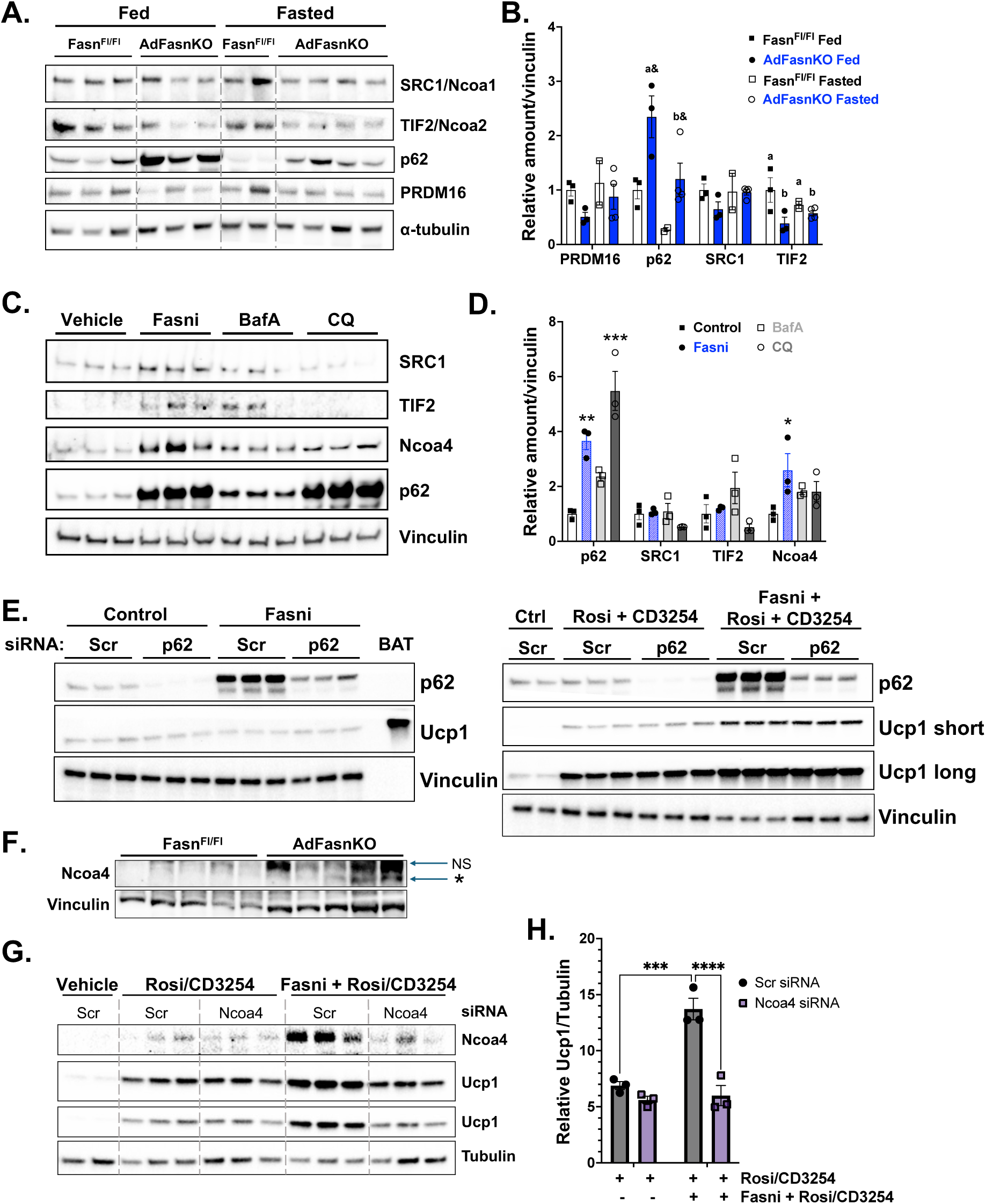
The PPARγ coactivator Ncoa4 is uniquely elevated in Fasn-deficient adipocytes. A) Fasn^Fl/Fl^ and AdFasnKO mice were fed or fasted for 24 hours. Western blot on subcutaneous WAT for PPARγ regulators and loading control, α-tubulin. B) Quantification of western blot in A. Proteins normalized to α-tubulin. 2-way ANOVA with Fisher’s LSD test. For p62: main effect of fasting p<0.05 and KO p<0.01, a& = p<0.05 vs. Fasn^Fl/Fl^ fed, b& = p<0.05 vs. AdFasnKO Fed; For TIF2: main effect of KO p<0.05, different letters are significantly different. C) *In vitro* differentiated primary adipocytes were treated with 100nM TVB-3664 (Fasni) for 48 hours or 100nM bafilomycin A (BafA) or 50uM chloroquine (CQ) for 24 hrs. Western blot for PPARγ regulators and loading control, Vinculin. D) Quantification of western blot in C. ANOVA with Dunnett post hoc *p<0.05, ** p<0.01, *** p<0.001 E) Western blot on subcutaneous WAT of Fasn^Fl/Fl^ and AdFasnKO mice for Ncoa4 with Vinculin as loading control. Asterisk points to Ncoa4, NS = nonspecific band. F) Scramble (Scr siRNA) or Ncoa4 knockdown (Ncoa4 siRNA) *in vitro* differentiated adipocytes were treated with 50nM rosiglitazone (Rosi) and 50nM CD3254 (CD) or DMSO with or without 100nM TVB-3664 (Fasni) for 48 hrs. Western blot for Ncoa4, Ucp1, and loading control, tubulin. G) Quantification of Ucp1 from western blot in F. 2-way ANOVA with Sidak’s multiple comparisons test *** p<0.001, **** p<0.0001.

Recently, the autophagy cargo receptor p62, which is consistently increased in FasnKO adipocytes and associated with autophagy inhibition, was shown to control PPARγ activity by competing with binding of the inhibitory NBR1 to the PPARγ:RXRα complex [25]. To determine whether increased p62 in FasnKO adipocytes mediates the effects on thermogenic gene expression, we utilized siRNA to silence p62 in mature adipocytes. Knockdown of p62 effectively eliminated the increase in p62 found with Fasn inhibition; however, agonist-induced Ucp1 protein was not affected (Figure 2e). Together, these data suggest the known PPARγ regulators Ncoa1/SRC1, Ncoa2/TIF2, PRDM16, and p62 do not mediate the effects of FasnKO on PPARγ target gene expression and thermogenesis.

### Ncoa4 is uniquely elevated in Fasn-deficient adipocytes

We next focused our attention on Ncoa4, Nuclear receptor coactivator 4, a protein related to Ncoa1 and Ncoa2. Like Ncoa1 and Ncoa2, Ncoa4 can function as a coactivator of PPARγ in cells with a PPRE reporter construct, in addition to other nuclear receptors including PPARα, AR, and others [26,27]. Importantly, in those studies, Ncoa4 increased ligand-activated PPARγ:RXRα activity [27]. Despite this, the function of Ncoa4 in adipose tissue remains unknown and unexplored. Notably, Ncoa4 also functions as the autophagy receptor for ferritin and mediates the process of ferritinophagy [28]. As a result, Ncoa4 protein levels are regulated in an autophagy-dependent manner, like p62, Ncoa1, and Ncoa2 [29]. We thus asked whether Ncoa4 is increased in FasnKO or Fasn-inhibited adipocytes and whether it could mediate the effects of Fasn deficiency on PPARγ activity. Indeed, Fasn inhibition in mature, fully differentiated adipocytes increased Ncoa4 protein levels (Figure 2c and 2d). Autophagy inhibition by Bafilomycin A or chloroquine also increased Ncoa4 levels, as expected, though not at statistically significant levels (Figure 2c and 2d). Importantly, Ncoa4 protein levels were increased in the adipose tissue of AdFasnKO mice (Figure 2f). Therefore, of the nuclear receptor coactivators investigated here (Ncoa1, Ncoa2, PRDM16), only Ncoa4 was upregulated in Fasn-deficient adipocytes and adipose tissue.

### Knockdown of Ncoa4 blocks the upregulation of Ucp1 by Fasn inhibition

To determine if the upregulation of the PPARγ coactivator Ncoa4 in response to Fasn inhibition mediates the effects on thermogenic gene expression, we used siRNA to silence Ncoa4 in mature differentiated adipocytes. Knockdown of Ncoa4 prevented the upregulation of Ncoa4 in response to Fasn inhibition (Figure 2g). Remarkably, knockdown of Ncoa4 completely blocked the PPARγ:RXRα agonist-induced upregulation of Ucp1 in response to Fasn inhibition (Figure 2g and 2h). Thus, in adipocytes *in vitro*, preventing the degradation of the PPARγ coactivator Ncoa4 via Fasn and autophagy inhibition is sufficient to drive PPARγ:RXRα-dependent Ucp1 expression.

### Ncoa4 enhances PPARγ target gene expression but not beiging of AdFasnKO adipose tissue

Based on the above, we asked whether Ncoa4 was also sufficient to drive the beiging of AdFasnKO white adipose tissue *in vivo*. Conditional Ncoa4^Fl/Fl^ mice were crossed with AdFasnKO (Fasn^Fl/Fl^; Adipo-Cre+) mice to generate adipocyte-specific Fasn and Ncoa4 double knockout mice (DKO; Fasn^Fl/Fl^; Ncoa4^Fl/Fl^; Adipo-Cre+). Surprisingly, analysis of thermogenic gene expression (Figure 3a) and morphological analysis of the subcutaneous adipose depot showed the DKO mice had similar levels of beiging to the single AdFasnKO mice (Figure 3b). Protein levels of Ucp1 in the subcutaneous depot were also similar between AdFasnKO and DKO mice (Figure 3c).

**Figure 3.**
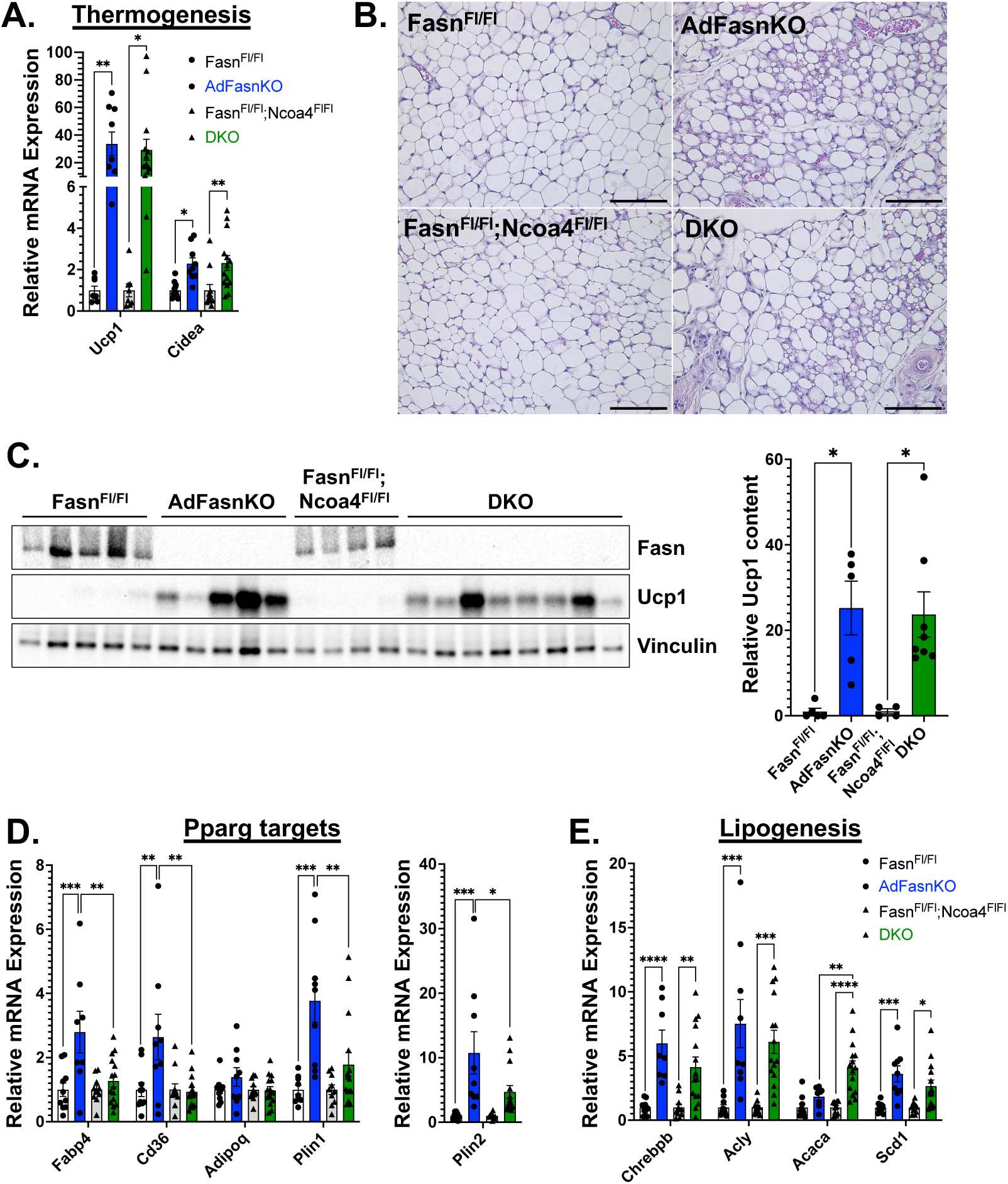
Double knockout of Ncoa4 and Fasn in iWAT prevents the upregulation of PPARγ genes by FasnKO alone but does not affect FasnKO-induced beiging. Subcutaneous WAT of Fasn^Fl/Fl^, AdFasnKO, Fasn^Fl/Fl^; Ncoa4^Fl/Fl^, and DKO mice A) qPCR analysis of thermogenic genes. 2-way ANOVA with Sidak’s multiple comparisons test * p<0.05, ** p<0.01 B) Hematoxylin and eosin staining of subcutaneous WAT. Scale bar = 100um. C) Western blot for Fasn, Ucp1, and Vinculin (loading control) and quantification of Ucp1 relative to Vinculin. 2-way ANOVA with Sidak’s multiple comparisons test * p<0.05 D) qPCR of “classic” PPARγ target genes. E. qPCR of genes in the lipogenesis pathway. 2-way ANOVA with Sidak’s multiple comparisons test * p<0.05, ** p<0.01, *** p<0.001, **** p<0.0001.

In addition to thermogenic genes, other PPARγ target genes are increased by FasnKO. Gene expression analysis of “classic” PPARγ target genes (Fabp4, Cd36, Plin1, Plin2) revealed that Ncoa4 was in fact required for their upregulation in AdFasnKO adipose tissue (Figure 3d). However, other PPARγ targets, including those in the lipogenic pathway, that were upregulated by FasnKO were unaffected in the double KO mice (Figure 3e). Genes in the lipogenic pathway are controlled not only by PPARγ, but also by the cAMP/PKA pathway and lipogenic transcription factors such as Chrebp and Srebp1c. Thus, it’s likely these factors override effects of Ncoa4 KO in the AdFasnKO adipose tissue. For instance, Chrebpb, the transcriptionally active isoform, is upregulated in AdFasnKO adipose tissue and remains upregulated in the DKO adipose tissue (Figure 3e), suggesting that Chrebpb, and/or other factors may drive lipogenic gene expression in these tissues. Similarly, the cAMP/PKA pathway is a major driver of thermogenic gene expression in adipose tissues, and our previous studies showed that this pathway was required for the beiging of AdFasnKO adipose tissue [20]. Therefore, we hypothesize that FasnKO adipocytes enhance cAMP/PKA signaling, which induces beiging, and which is independent of Ncoa4 upregulation. However, Ncoa4 is absolutely required for the activation of classic PPARγ target genes in AdFasnKO subcutaneous white adipose tissue.

### Ncoa4 is required for adipogenesis in wild-type adipocytes

The data in Figures 1-3 suggested that while not required for FasnKO-induced beiging, Ncoa4 can control PPARγ-dependent gene expression in adipocytes. As PPARγ is a major regulator of adipogenesis, we asked whether Ncoa4 contributes to adipogenesis of isolated primary preadipocytes. We first analyzed the expression profile of Ncoa4 during the differentiation of adipocytes *in vitro*. Shown in Figure 4a, Ncoa4 expression is lowest in preadipocytes (Day 0) prior to differentiation, spikes at the onset of differentiation, and remains elevated in mature adipocytes (Day 5). Interestingly, the spike in Ncoa4 expression coincides with the onset of Fabp4 expression, an adipogenic marker and PPARγ-regulated gene, and immediately precedes major spikes in Adipoq expression. Protein levels of Ncoa4 mirror the changes in RNA expression during differentiation (Figure 4b). Crispr/Cas9 gene editing to delete Ncoa4 was then performed in mouse primary preadipocytes. Two different guide RNAs were used and showed different levels of Ncoa4 knockout, illustrated by the amount of Ncoa4 mRNA in Figure 4c. Strikingly, Ncoa4 KO in preadipocytes strongly impairs differentiation and lipid accumulation (Figure 4d), the extent to which correlates with the degree of Ncoa4 suppression (Figure 4c).

**Figure 4.**
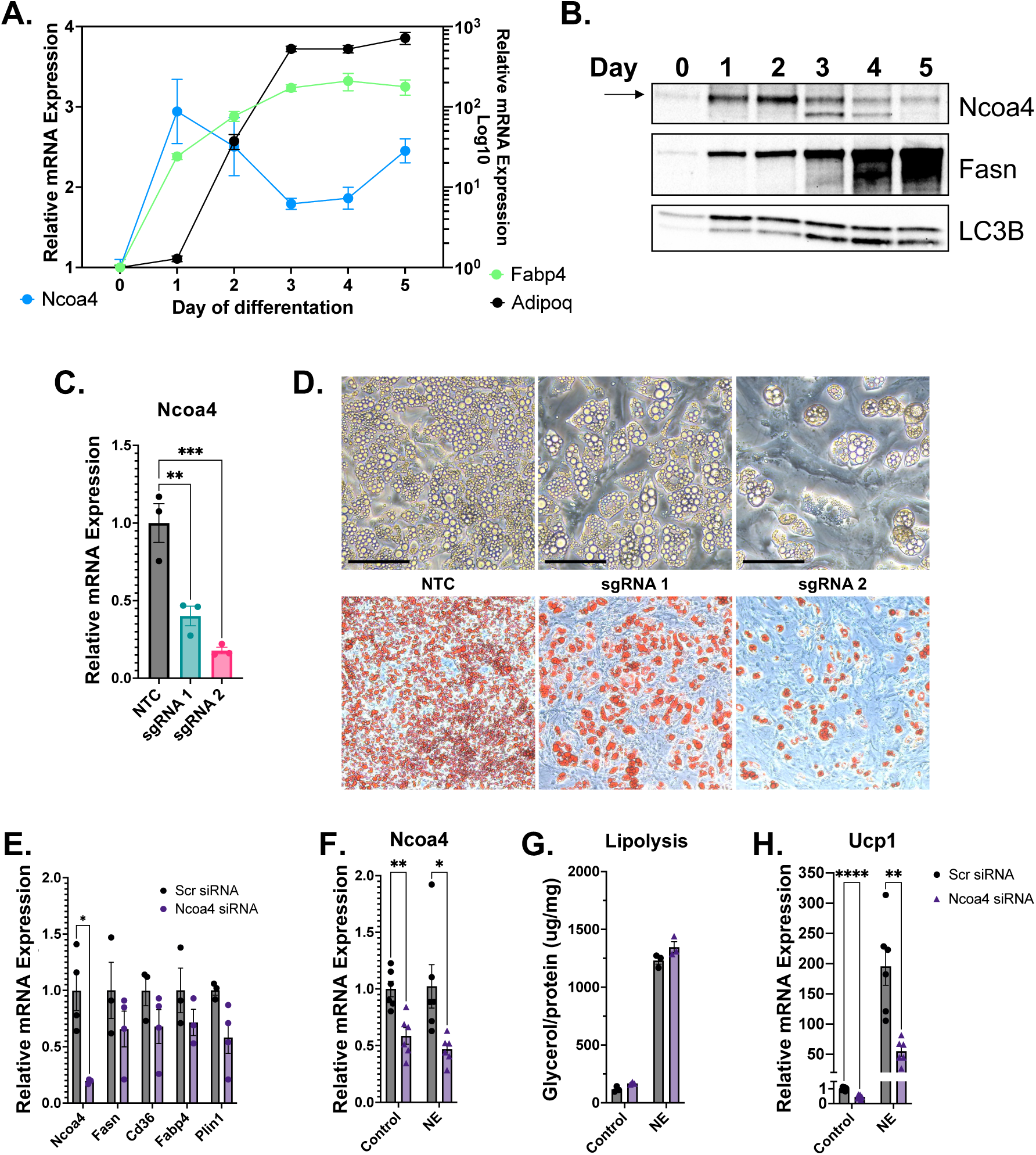
Loss of Ncoa4 impairs adipogenesis and thermogenic gene expression in cultured white adipocytes. Time course of Ncoa4, Fabp4, and Adipoq mRNA expression during the differentiation of primary adipocytes differentiated *in vitro*. Relative Ncoa4 expression is on the left axis, and relative Fabp4 and Adipoq expression are on the right axis in logarithmic scale. B. Western blot for Ncoa4, Fasn, and LC3B during the differentiation of primary adipocytes. C. Crispr/Cas9-edited mouse primary adipocytes were differentiated *in vitro* using 1 of 2 different gRNAs targeting Ncoa4 (sgRNA1 or sgRNA2) or unedited (NTC). qPCR analysis of Ncoa4 mRNA expression in differentiated edited adipocytes. ANOVA with Tukey’s post hoc ** p<0.01, *** p<0.001 D) Light microscopy images (top panels) and oil red O staining (bottom panels) of Crispr-edited differentiated adipocytes. Scale bar = 50um. E) After differentiation, mature *in vitro* differentiated adipocytes were transfected with scramble (Scr) or Ncoa4 siRNA. qPCR analysis of adipogenic genes in Scr or Ncoa4 knockdown (Ncoa4 siRNA) adipocytes. Student’s t test * p<0.05 F) Mature adipocytes were treated as in E. 48 hrs post-transfection adipocytes were stimulated with 100nM norepinephrine (NE) or vehicle (control) for 3 hrs. qPCR analysis of Ncoa4 expression after NE stimulation. G) Glycerol release into media from adipocytes in F. H) qPCR analysis of Ucp1 expression in adipocytes in F. 2-way ANOVA with Sidak’s multiple comparisons test * p<0.05, ** p<0.01, **** p<0.0001 (for F,G,H)

### Ncoa4 is required for maximal norepinephrine-stimulated Ucp1 expression but not lipolysis in mature adipocytes

To determine the role of Ncoa4 in mature adipocytes, we utilized siRNA to silence Ncoa4 after differentiation. In unstimulated conditions, Ncoa4 knockdown (KD) did not affect adipogenic markers (Figure 4e), lipid content (data not shown), or norepinephrine (NE)-stimulated lipolysis (Figure 4f and 4g). However, when acutely stimulated with NE, Ncoa4 knockdown significantly blunted Ucp1 induction (Figure 4f and 4h), suggesting Ncoa4 may contribute to thermogenesis.

Multiple single nuclei RNA sequencing datasets investigating the effects of cold exposure on brown and white adipocytes are now publicly available [30,31]. We took advantage of these datasets to query whether Ncoa4 expression correlates with thermogenic gene expression *in vivo*. In white adipocytes [31], Ncoa4 expression increased after 4 and 7 days of cold exposure (Supplemental Figure 1a) and largely overlapped with adipocyte populations expressing Ucp1 (P2 and P5), though some subclusters express Ncoa4 and Ucp1 exclusively (P3 and P4) (Supplemental Figure 1b). Likewise, in brown adipocytes [30], Ncoa4 was more highly expressed after cold exposure (Supplemental Figure 1c, reflected in populations P6-P10), though in contrast to white adipocytes, Ncoa4 and Ucp1 expression share a more similar distribution pattern across subclusters (Supplemental Figure 1c). Collectively, these data indicate Ncoa4 plays an important role in adipocyte differentiation and may also play an important role in mature adipocyte functions, including thermogenesis.

### AdNcoa4KO mice exhibit impaired brown fat response to cold exposure

To understand the role of Ncoa4 in adipose tissue *in vivo*, we used the adipocyte-specific Ncoa4 knockout (AdNcoa4KO) mice. In chow-fed, room temperature-housed mice, AdNcoa4KO did not affect adipogenic, lipogenic, or thermogenic gene expression in subcutaneous WAT (Figure 5a) or BAT (Figure 5b). Body weights and adipose tissue weights were similarly unaffected (Supplemental Figure 2a). Since our *in vitro* studies showed that Ncoa4 KD suppresses Ucp1 expression and Ncoa4 is induced by cold exposure, we asked whether Ncoa4 is required for thermogenesis in vivo. To this end, AdNcoa4KO mice were challenged with an acute cold exposure. During the acute phase, AdNcoa4KO mice were cold-tolerant; however, during prolonged cold exposure, AdNcoa4KO mice were unable to defend body temperature as well as controls (Figure 5c). After 1 week of cold exposure, body weights and adipose tissue weights did not differ between Ncoa4^Fl/Fl^ and AdNcoa4KO mice (Supplemental Figure 2b). Interestingly, Ncoa4 protein levels were increased in the subcutaneous WAT of cold-exposed Ncoa4^Fl/Fl^ mice (Figure 5d). Surprisingly, however, cold-induced beiging of the WAT was similar between Ncoa4^Fl/Fl^ and AdNcoa4KO mice (Figure 5d and 5e).

**Figure 5.**
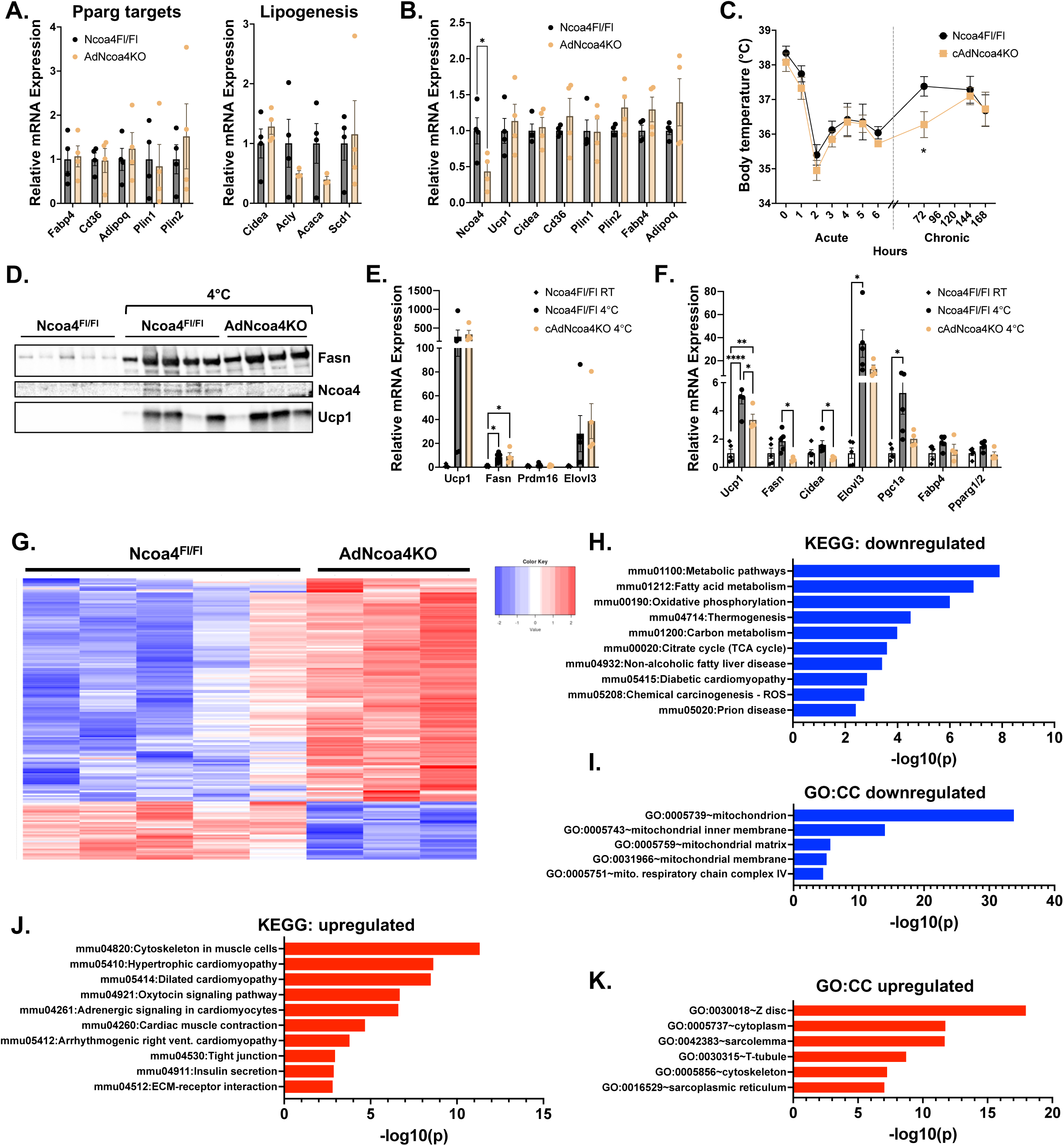
AdNcoa4KO mice have impaired BAT thermogenesis in response to cold. A) qPCR analysis of PPARγ target genes in subcutaneous WAT from Ncoa4^Fl/Fl^ and AdNcoa4KO mice. B) qPCR analysis of PPARγ target genes in BAT from Ncoa4^Fl/Fl^ and AdNcoa4KO mice. Student’s t test * p<0.05 C) Rectal temperature of Ncoa4^Fl/Fl^ and AdNcoa4KO mice during an acute cold challenge (6°C). Student’s t test * p<0.05 D) Western blot analysis of Fasn, Ncoa4, and Ucp1 in the subcutaneous WAT of 1 week cold-exposed Ncoa4^Fl/Fl^ and AdNcoa4KO mice and room temperature-housed Ncoa4^Fl/Fl^ mice. E) qPCR analysis of subcutaneous WAT from mice as in D. ANOVA with Tukey’s post hoc * p<0.05 F) qPCR analysis of PPARγ targets and thermogenic genes in BAT from mice as in D. ANOVA with Tukey’s post hoc * p<0.05, ** p<0.01, **** p<0.0001 G) RNA sequencing was performed on BAT from 1 week cold-exposed Ncoa4^Fl/Fl^ and AdNcoa4KO mice. Heatmap showing differentially expressed genes with padj < 0.05. H) Kyoto Encyclopedia of Genes and Genomes (KEGG) pathway analysis of genes downregulated (padj < 0.1) in cold-exposed AdNcoa4KO BAT. I) Gene ontology (GO): cellular components (CC) analysis of downregulated genes (padj < 0.1) in cold-exposed AdNcoa4KO BAT. J) KEGG pathway analysis and K) GO: CC analysis of upregulated genes (padj < 0.1) in cold-exposed AdNcoa4KO BAT.

Brown adipose tissue is the major thermogenic organ in mice; therefore, the BAT of the cold-exposed AdNcoa4KO mice was assessed. Strikingly, the majority of cold-induced thermogenic genes examined were significantly reduced by AdNcoa4KO (Figure 5f). To probe these effects in greater detail, we performed bulk RNA sequencing on the cold-exposed BAT from Ncoa4^Fl/Fl^ and AdNcoa4KO mice. Analysis of the differentially expressed genes showed a large number of genes downregulated by AdNcoa4KO (Figure 5g). KEGG pathway analysis of these genes showed that the top pathways affected were involved in fatty acid metabolism, oxidative phosphorylation, and thermogenesis (Figure 5h). In addition, GO cellular components analysis showed that genes involved in mitochondrial function were significantly reduced by Ncoa4KO (Figure 5i).

To our surprise, a greater number of genes were upregulated in the cold-exposed AdNcoa4KO BAT (Figure 5g). KEGG pathway analysis and GO cellular components analysis of these upregulated genes showed a striking and consistent trend of pathways involved in skeletal and cardiac muscle function (Figure 5j and 5k). Together these data suggest Ncoa4 is required for optimal thermogenesis in BAT, and loss of Ncoa4 reduces thermogenic and mitochondrial gene expression, while inducing a myogenic gene signature.

## Discussion

PPARγ is a major driving regulator of white and brown adipose tissue biology [3]. Identifying mechanisms that promote PPARγ activity can therefore have great implications for metabolic health. Recent studies have shown that autophagy can control PPARγ activity through the turnover of PPARγ coregulators [11,12], and the studies described here provide further evidence for that notion. We show that autophagy deficiency is associated with an increased availability of a PPARγ coactivator, Ncoa4, and increased PPARγ target gene expression. We also demonstrate that Ncoa4 is a critical player in brown adipose tissue thermogenesis, and together, the data presented here firmly establish Ncoa4 as a regulator of adipocyte PPARγ activity and white and brown adipose tissue function.

The mechanism underlying the unexpected beiging of AdFasnKO adipose tissue has remained elusive. We previously found that both macrophages and the cAMP/PKA pathway appear required optimally for this beiging; however, the cell autonomous changes within FasnKO adipocytes that elicit macrophage recruitment or polarization and/or affect cAMP/PKA signaling are still unknown [20]. Our finding that PPARγ activity is potentiated in FasnKO adipocytes in a cell-autonomous manner led us to hypothesize this could be a mechanism whereby FasnKO adipocytes are sensitized to beiging stimuli, since PPARγ is required for full activation of the thermogenic program. Indeed, we show that preventing Ncoa4 degradation by FasnKO-induced autophagy impairment activates PPARγ activity. However, our data here definitively show that preventing PPARγ activation in FasnKO adipocytes in vivo by Ncoa4KO does not block beiging of the FasnKO adipose tissue *in vivo*. These data suggest another factor is involved in the beiging of FasnKO adipose tissue that is independent of Ncoa4-driven PPARγ activity and likely modulates macrophage recruitment or PKA signaling.

In an earlier study, Lodhi et al. concluded FasnKO adipocytes exhibit decreased PPARγ activity, due to reduced ligand synthesis, and a shift toward PPARα activity, which is in seeming contrast to our data [18]. However, certain points suggest these studies may in fact have common, non-contrasting conclusions. In addition to PPARγ, PPARα was also a top hit in our transcription factor analysis for the FasnKO adipose tissue RNASeq data (Figure 1a), explained in part by the fact that PPARγ and PPARα share many common target genes. Another possible explanation is that nuclear receptor coregulators, including Ncoa4, Ncoa1/SRC1, and Ncoa2/TIF2 are promiscuous and can regulate various nuclear receptors, including PPARγ and PPARα. These data suggest that both PPARγ and PPARα may be more active in FasnKO adipocytes, as previously reported. However, in contrast to the previous report, our data show that in the presence of specific agonists for PPARγ and RXRα, PPARγ-dependent gene expression is potentiated by FasnKO. The study by Lodhi et al. did not examine the effects of FasnKO with agonists for both PPARγ and RXRα, which may explain the discrepancy. These data may also explain why FasnKO adipocytes *in vitro* require agonist stimulation for beiging/PPARγ activation, while in vivo, FasnKO adipocytes do not require exogenous agonist stimulation. Due to the presence of multiple non-adipocyte cell types in adipose tissue *in vivo* and constant exposure to circulating factors, it’s likely the FasnKO adipocytes are exposed to ligands for PPARγ, RXRα, and other nuclear receptors. However, in the closed *in vitro* system, these ligands are absent or at low levels and need to be supplemented to replicate the in vivo setting.

In addition to functioning as a nuclear receptor coactivator, it’s well established that Ncoa4 serves as a receptor for ferritin in the process of ferritinophagy [28]. When cells are exposed to low intracellular iron levels, Ncoa4 binds the iron-storage protein ferritin and delivers it to the lysosome, where free iron can be released [28]. As a result, Ncoa4 availability regulates intracellular iron levels [32]. In various cell types, loss of Ncoa4 has been shown to reduce intracellular iron availability and thus has been implicated in processes such as ferroptosis and erythropoiesis and can affect enzymatic activities involved in epigenetic modifications, mitochondrial function, and other functions [29,33–36]. The studies presented here have not examined the role of Ncoa4 in adipocyte iron homeostasis. Such studies are the focus of current and future investigations and will delineate whether Ncoa4KO-induced iron alterations (if any) contribute to the phenotypes of AdNcoa4KO adipocytes and mice.

Our data show that Ncoa4 is required for optimal cold-induced thermogenesis in BAT in mice. We also found that Ncoa4 expression is increased by cold exposure in BAT and subcutaneous WAT. Using previously published single nuclei sequencing datasets, we confirmed that Ncoa4 is increased specifically in brown and white adipocytes after cold exposure, and its expression is most associated with thermogenic adipocytes, i.e. those expressing Ucp1, underscoring its importance for thermogenesis. PPARγ is required for mature brown adipocyte thermogenesis and inducibility by b-adrenergic stimuli, including cold exposure [4]; therefore, we hypothesize that Ncoa4 enables BAT thermogenesis by enhancing PPARγ activity through its coactivator function. However, we cannot rule out the contribution of Ncoa4 to intracellular iron availability and potential effects on thermogenesis, which are the focus of current studies.

One striking effect of AdNcoa4KO on cold-exposed BAT was the induction of a myogenic gene signature. In mice, BAT and skeletal muscle share a common developmental precursor, and fate determination into brown adipocytes is controlled by the coregulatory protein PRDM16, which regulates transcription factors including PPARγ, Pgc1a, and C/EBP-b [37]. Interestingly though, ablation of PRDM16 or PPARγ in brown adipocytes does not induce a myogenic signature, suggesting that a loss of Ncoa4 may not be simply impairing PPARγ or PRDM16/PPARγ complex transcriptional activity [4,38,39]. Instead, loss of epigenetic modifiers, including the histone methyltransferase, Ehmt1, or the histone demethylase, UTX, can induce a myogenic signature in BAT [40,41]. These data suggest Ncoa4 may affect the activities of epigenetic modifier enzymes or may affect their binding/recruitment to regulatory complexes, including the PRDM16/PPARγ complex.

Lastly, in the subcutaneous WAT of cold-exposed mice, Ncoa4 is upregulated, yet the loss of Ncoa4 has no effect on the beiging of WAT. This contrasts with our *in vitro* data, which showed that Ncoa4KD impairs norepinephrine-induced upregulation of Ucp1. A plausible explanation is that the defective BAT thermogenesis observed in AdNcoa4KO mice induces a compensatory increase in white adipose beiging and thus masks the effect of AdNcoa4KO. Future studies will parse the effects of cold exposure on white vs. brown adipocytes *in vivo*.

In conclusion, these studies have revealed that Ncoa4 is a novel regulator of brown and white adipose tissue biology, hypothetically through direct control of PPARγ activity. Ncoa4 levels can be modulated by autophagy, suggesting autophagy is a major contributor to mature adipocyte functions. Powerful signaling nodes in adipose tissue, including the insulin pathway and mTORC1, directly control autophagy flux in adipocytes, suggesting these pathways could affect adipocyte health in part via control of Ncoa4 levels. Modulators of autophagy and/or Ncoa4 are thus potentially interesting therapeutic agents to control adipose tissue thermogenesis, PPARγ activity, and metabolic health.

## Methods

### Animals

All animal procedures were approved by the University of Massachusetts Chan Medical School Institutional Animal Care and Use Committee (IACUC). Mice were housed in ventilated cages with automated watering systems, on a 12h-light/12h-dark light cycle, at 21-23°C, and 30-35% humidity. Mice were fed normal chow diet (LabDiet, 5P76 Prolab® RMH 3000) ad libitum with free access to water. Adipocyte specific-Fasn KO mice (AdFasnKO) were generated by crossing Fasn^Fl/Fl^ with Adiponectin-Cre mice [42,43] (obtained from Jackson Labs (Jackson Labs, 028020). Adipocyte specific-Ncoa4 KO mice (AdNcoa4KO) were generated by crossing Ncoa4^Fl/Fl^ (Jackson Labs, 033295) with Adiponectin-Cre mice. Wildtype (C57Bl/6J) male and female mice were bred in house and used at 3-5 weeks of age to prepare primary adipocyte cultures, unless knockout strain is noted. Animal tissues harvested for protein and RNA consisted of 10–12-week-old male mice on a normal chow diet.

### Cold Exposure

AdNcoa4KO and Ncoa4^Fl/Fl^ mice were challenged with cold temperature by placing mice in cages (singly housed) with free access to water in an environmental chamber set at 6°C. During the first 6 hours, the mice were not given food but were given free access to food for the remaining days of the cold exposure. Core body temperature was monitored hourly during the first 6 hours, followed by day 3, day 6 and day 7. Temperature was measured by using a rectal probe (Braintree Scientific, RET-3). On day 7, animals were euthanized and tissues harvested for downstream analysis.

### Primary Preadipocyte Isolation, Maintenance, and Adipocyte Induction

Male and female mice were euthanized and inguinal subcutaneous fat pads were harvested for primary cell culture. Briefly, subcutaneous fat pads were harvested with the lymph node intact and placed in Hanks’ Balanced Salt Solution (Invitrogen, 14025-092) containing 2% bovine serum albumin (Sigma, A9647). Tissue was minced to 2-5mm pieces with scissors, then digested by adding collagenase (Sigma, #C6885) for a final concentration of 1.5 mg/ml for 30-45 min at 37°C, with gentle shaking. Fetal bovine serum was added to the digestion mix to neutralize the collagenase, and the cell suspension was filtered with a 100um cell strainer and centrifuged at 500xg to pellet the stromal vascular fraction (SVF). The SVF was resuspended in red blood cell lysis buffer (Invitrogen, 00-4333-57), kept at room temperature for 2-3 min, then centrifuged, and the pellet was resuspended in growth media consisting of DMEM-F12 (Gibco,11330-032), 10% Fetal Bovine Serum (R&D Systems, S11550), 1% Penicillin-Streptomycin (Gibco, 151140122) and 100 ug/ml Normocin (InvivoGen, ant-nr-1), followed by filtering through a 40um cell strainer. The final filtered cell suspension was plated and cells were maintained in a cell incubator with 5% CO2, at 37 °C. To induce adipocyte differentiation, cells were grown to confluence, then cultured with induction media consisting of growth media listed above supplemented with: 5 μg/mL insulin, 1 μM dexamethasone, 0.5 mM 3-isobutyl-1-methylxanthine, 60 μM indomethacin and 1 μM rosiglitazone. After 48 hrs, the media was switched to growth media supplemented with 5 μg/mL insulin, and cells were fed with the same media on day 4. On day 5, the cells were considered fully differentiated.

For rosiglitazone and CD3254 treatments on wild-type cells, adipocytes were treated on day 4 or day 5 with 50 nM rosiglitazone (Cayman Chemical, 11884) and 50 nM CD3254 (Cayman Chemical, 20870) for 48 hrs. Concurrently, the Fasn inhibitor TVB3664 (Selleck Chemical, S8563) was added at 100nM. Vehicle-treated cells were given DMSO at the same concentration. For autophagy inhibition, 100nM bafilomycin A (Cayman Chemical, 11038) or 50μM chloroquine (Sigma, C6628) were added to cells for 24 hrs.

For AdFasnKO primary adipocytes, 50nM rosiglitazone and 50nM CD3254 were administered on day 3 of differentiation. Adipocytes were cultured for 72 hrs and harvested on day 6.

### Genetic Editing of Primary Adipocytes

Primary preadipocyte cell cultures were transfected with 2 different guide RNAs to Ncoa4 using the 100ul Neon Transfection Kit (Invitrogen, MPK10025). Briefly, preadipocytes were grown as described above. CRISPR ribonucleoprotein (RNP) complexes were prepared by diluting SpyCas9 enzyme (PNA Bio, #CP02) or 3xNLS-SpCas9 (purified by the Scot Wolfe lab) and sgRNA (IDT, sgRNA #1 5’ CACGCGAGCTCCTCAAGTAT 3’ sgRNA #2 5’ GCAGGCTCAGCAGCTCTATT 3’) with Buffer R to deliver 300pmol Cas9 and 400pmol sgRNA per 2.5×10^6^ cells. The complexes were incubated at room temperature for 20min. Cells were prepared while RNP complexes formed by trypsinizing, washing with PBS, and resuspending in buffer R. For each electroporation 60ul of RNP complexes was mixed with 60ul of cells in Buffer R for a final amount of 2.5×10^6^ cells. Cells were electroporated in the Neon Electroporation Machine with the settings, 1350V-30ms-1 pulse. Immediately after electroporating, cells were plated in growth media and grown and differentiated as described above. Genetic editing was confirmed by PCR amplification of DNA harvested from cells post-transfection. After Sanger sequencing (Genewiz), amplicons were fed into the Synthego ICE Crispr Analysis Tool. Knockouts were further validated by qPCR and western blot analysis.

### siRNA Treatment of Primary Adipocytes

Primary adipocytes from wild-type mice were isolated, grown, and differentiated. On day 4-5 of differentiation, cells were treated with 50nM Scr siRNA-A (Santa Cruz, sc-37007) or Ncoa4 siRNA (Santa Cruz, sc-29720) in RNAiMAX reagent (Invitrogen, 13778-150) according to the manufacturer’s instructions. Briefly, siRNA-RNAiMAX complexes were prepared in Opti-MEM, diluted with growth media, and added to the adipocytes. 48-72hrs after transfection, the knockdown was validated by RT-PCR.

### Glycerol Release Assay

Primary adipocytes were transfected with siRNA as described above. Approximately 48 hours after siRNA introduction, cells were washed with PBS and glucose- and pyruvate-free DMEM (Gibco #14430-01) supplemented with 4% BSA was added along with 100nM norepinephrine for 2-3hrs or vehicle. Media was collected for use in free glycerol determination kit (Sigma, FG0100) following the manufacturer’s instructions. Glycerol content was normalized to protein concentration.

### Histology and Oil Red O Staining

Subcutaneous fat tissue was fixed in 10% neutral buffered formalin followed by embedding in paraffin. Tissue blocks were sectioned and stained with hematoxylin and eosin by the University of Massachusetts Chan Medical School Morphology Core.

Cells in culture dishes were stained with Oil Red O (ORO). Briefly, cells were fixed in 4% paraformaldehyde for 15 min at room temperature. Cells were then incubated in 60% isopropanol for 2 min. After 2 min, the isopropanol was removed, and ORO solution (Sigma, O1391) was added for 20 min at room temperature. ORO was removed, and the cells were washed with water. Images of H&E sections and ORO were visualized and imaged with a Leica DM2500 LED microscope equipped with a Leica MC170 HD camera with a 5x/0.12 objective or a 20x/0.4 Leica objective.

### Protein Isolation and Western Blotting

Whole pieces of fat tissue were homogenized in isolation buffer consisting of 50mM Tris-HCl (pH 8.0), 0.25M NaCl, and 5mM EDTA and 1x HALT protease inhibitors (Thermo Scientific, 78430). After homogenizing, the tissue was centrifuged at 6,000xg for 15 min at 4°C. The fat cake was removed and the infranatant and cell pellet were resuspended in the remaining buffer. SDS and Triton X were added to final concentrations of: 2% SDS and 1% Triton X-100, followed by incubation on ice for 30 min. Cell lysates were centrifuged for 15 min at 12,000 x g at 4°C and supernant was collected for analysis. Cultured cells were lysed by scraping on ice into isolation buffer listed above (including 1x HALT protease inhibitors) with 2% SDS and 1% Triton X-100 followed by vortexing and heating at 95°C for 5 min and sonication. Protein quantitation was performed using a Pierce BCA kit following manufacturer’s instructions (Pierce, A65453). Samples were prepared for western blotting by diluting the lysates using Laemmli buffer with 2-mercaptoethanol. Samples were heated at 95 °C for 10 min, followed by separation by SDS-PAGE using Mini-PROTEAN TGX Precast Protein Gels (Biorad). Proteins were transferred to nitrocellulose by using the Turbo-Blot mini nitrocellulose packs (BioRad, 1704158) and the Trans-Blot Turbo Mini System. Protein transfer was verified by incubating membranes with 0.1% Ponceau S solution (0.1% Ponceau S, 5% v/v glacial acetic acid in water) for 3 min with shaking, followed by rinsing with water. Blots were blocked in blocking buffer consisting of 5% BSA in 0.1% TBS-T (20mM Tris pH7.5, 150mM NaCl, 0.1% Tween-20) for 30-60 min at room temperature. Primary antibodies were incubated overnight at 4°C. Blots were incubated with secondary antibodies (1:10,000 goat anti-rabbit or -mouse IgG-HRP) at room temperature for 1 hr. Blots were developed with enhanced chemiluminescence substrate (Perkin Elmer, NEL1404001EA) following the manufacturer’s instructions, and imaged using a BioRad Chemidoc.

Proteins were blotted with the following antibodies: anti-Fasn (CST #3180), anti-Fabp4 (R&D #AF1443), anti-IgG LC (goat anti-mouse Invitrogen #31430), anti-p62 (CST #23214), anti-Ucp1 (Abcam #10983), anti-LC3B (CST #83506), anti-alpha-tubulin (Sigma #T5168), anti-SRC1/Ncoa1(CST #2191), anti-TIF2/Ncoa2(CST #96687), anti-PRDM16 (Abcam #ab106410), anti-Ncoa4 (Santa Cruz #sc-373739 or custom antibody made by Genscript), anti-vinculin (CST #18799)

### RNA Isolation and Gene Expression Analysis

RNA was isolated from cells in culture using Trizol reagent (Invitrogen, 15596026) according to the manufacturer’s instructions. RNA was isolated from tissue by taking whole frozen tissue and crushing in liquid nitrogen to produce tissue powder. Trizol was added to the powdered tissue with a stainless steel bead and homogenized using a Qiagen Tissue Lyzer II. For fat tissue, Trizol/tissue homogenate was centrifuged at 4°C, 12,000xg for 10 min to float the lipid to the surface. Trizol/tissue mixture was transferred to a new tube avoiding the lipid layer at the surface. RNA was isolated from the tissue following the manufacturer’s instructions. RNA concentrations were measured using a NanoDrop 2000. For gene expression analysis, 1ug of isolated RNA was reverse transcribed using BioRad iScript cDNA synthesis kit (BioRad, 15596026). Quantitative PCR (qPCR) was performed by using iTaq Universal SYBR Green Supermix (BioRad, 15596026) on a BioRad CFX96 System. Gene expression was quantified using the ΔΔCT method utilizing the following housekeeping genes as internal control reference genes: 36b4 and 18s. Primer sequences listed in Table 1.

**Table 1.**
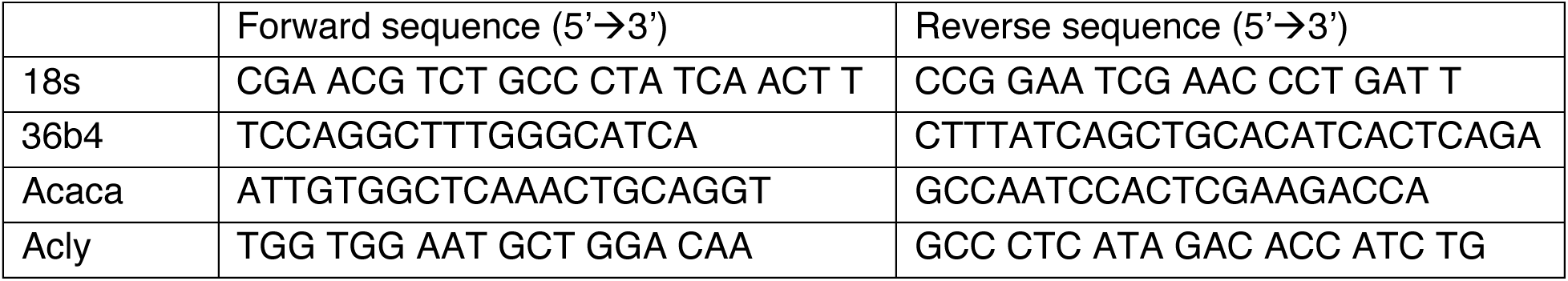
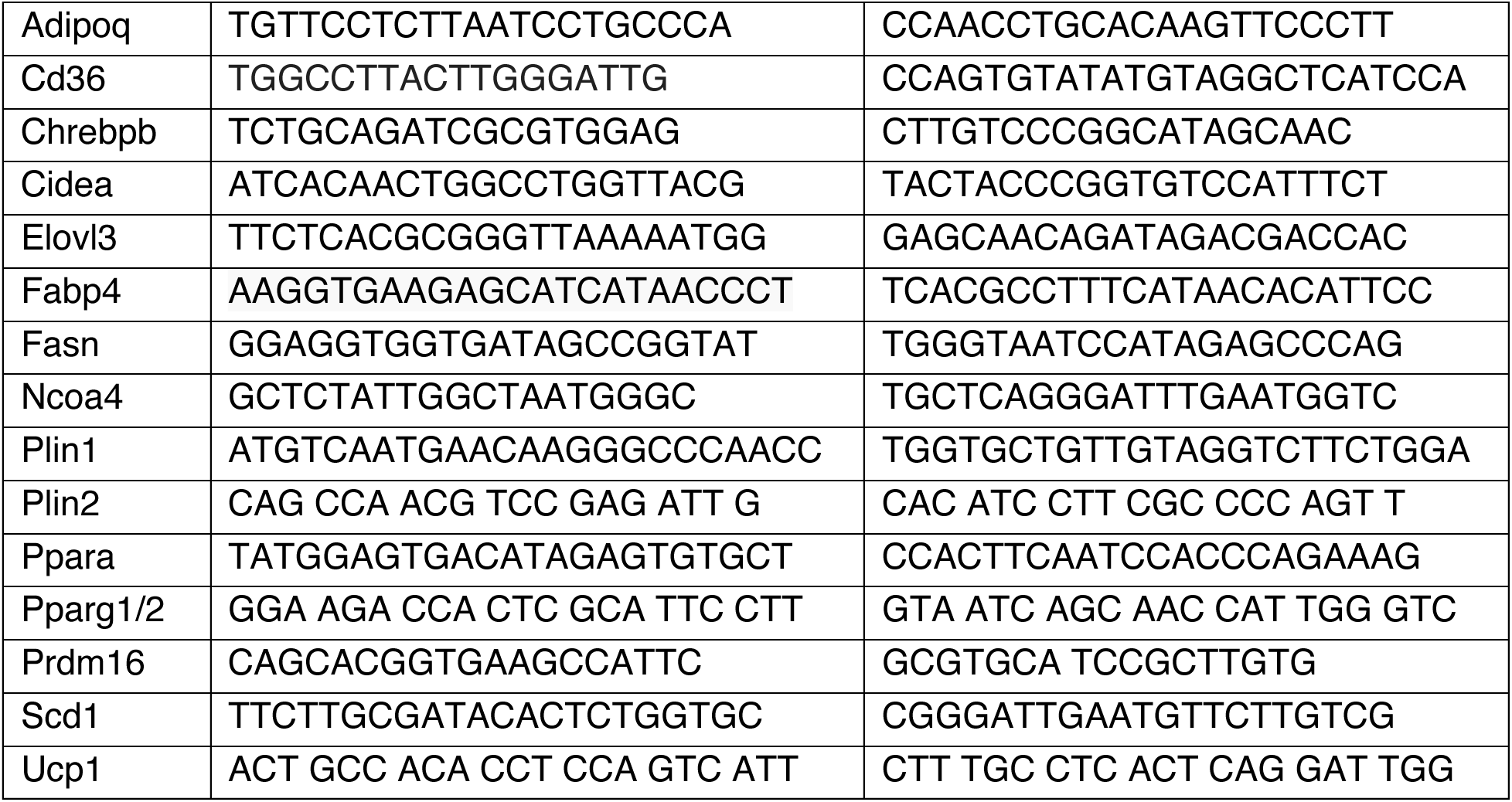
Primer sequences for qPCR.

### RNA Sequencing

Total RNA was isolated as described above from the subcutaneous white adipose tissue from 1-week cold-exposed AdNcoa4KO and Ncoa4^Fl/Fl^ mice and submitted to Azenta/Genewiz for standard RNA sequencing (Illumina 2×150bp) with DNase treatment and rRNA depletion. Demultiplexed FASTQ files were obtained and entered into the DolphinNext RNA-Seq pipeline [44] for sequence alignment and quantification of gene expression with the following parameters: 3’ adapter removal (Trimmomatic V0.32), Sequential Mapping, STAR 2.7.11b and RSEM for alignments, and FastQC for quality control. Differential gene expression analysis was performed using DEBrowser [45], and data were normalized using the RLE method. The heatmap was generated in DEBrowser showing differentially expressed genes with padj < 0.05 after normalization. Pathway and GO term analyses were performed using DAVID [46,47]. Data entered were up or downregulated genes (FC>1 or <1, respectively, with padj<0.1). TRUSST analysis [22] was performed using Metascape [48].

## Acknowledgements

We thank all members of the Czech laboratory for helpful discussions and feedback. Funding was also provided by grants from the National Institute for Diabetes and Digestive and Kidney Diseases (DK116056 and DK030898 to M.P.C.). We thank the Scot Wolfe lab at UMass Chan Medical School for providing purified Cas9 enzyme.

## Author Contributions

L.A.R. and M.P.C. conceived the study and designed the experiments. L.A.R., K.D.S., A.G., S.M., M.Y. performed the experiments. L.L. and L.A.R. performed single cell data analysis, and L.A.R. analyzed the bulk RNA sequencing data. S.N. contributed to the methods section, and S.N. and H.W. helped with mouse tissue collection. L.A.R. and M.P.C. wrote the manuscript, which was reviewed and edited by all co-authors.

---

**Supplemental Figure 1.**
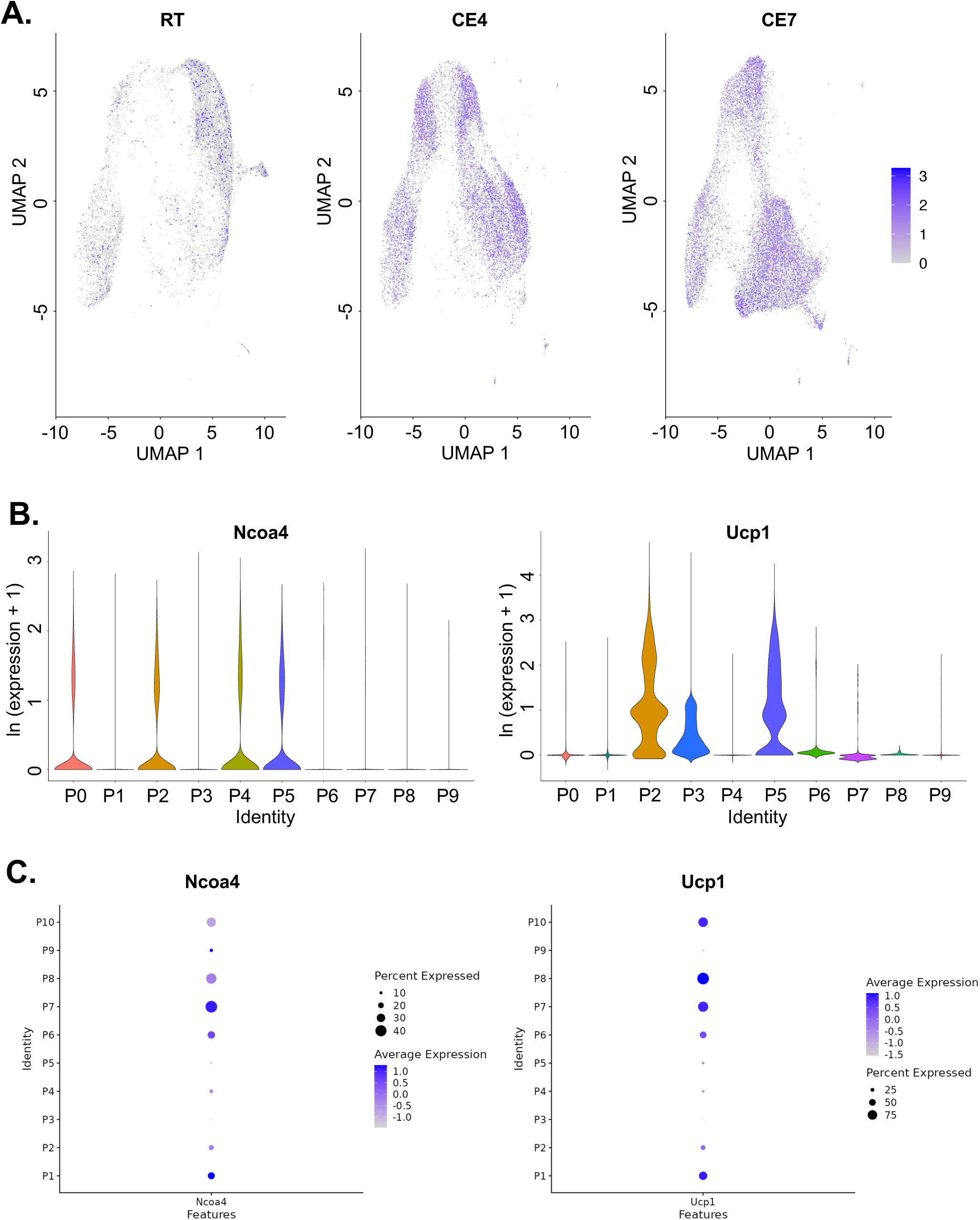
Ncoa4 is associated with thermogenic adipocyte populations in single nuclei sequencing datasets. A) Data from [31]. Seurat Feature plot of Ncoa4 from its “RNA” assay “data” slot, i.e. of ln of sample depth normalized expression counts. Data is split into room temperature (RT), 4-days cold exposure (CE4) and 7-days cold exposure (CE7). B) Ucp1: Seurat VlnPlot of ln of depth normalized and “integrated” expression data. The widths are all normalized to have the same maximum value (the default for VlnPlot). Ncoa4: similarly but prior to integration. C) Data downloaded from batnetwork.org, from [30]. Dot plots show mouse Ncoa4 and Ucp1 expression across brown adipocyte subclusters. Clusters P6-P10 are mainly associated with cold-exposed adipocytes, P1 and P2 from room-temperature and cold-exposure groups, P3 from the thermoneutral group, P5 from the room-temperature group, and P4 reflects adipocytes from all temperature groups.

**Supplemental Figure 2.**
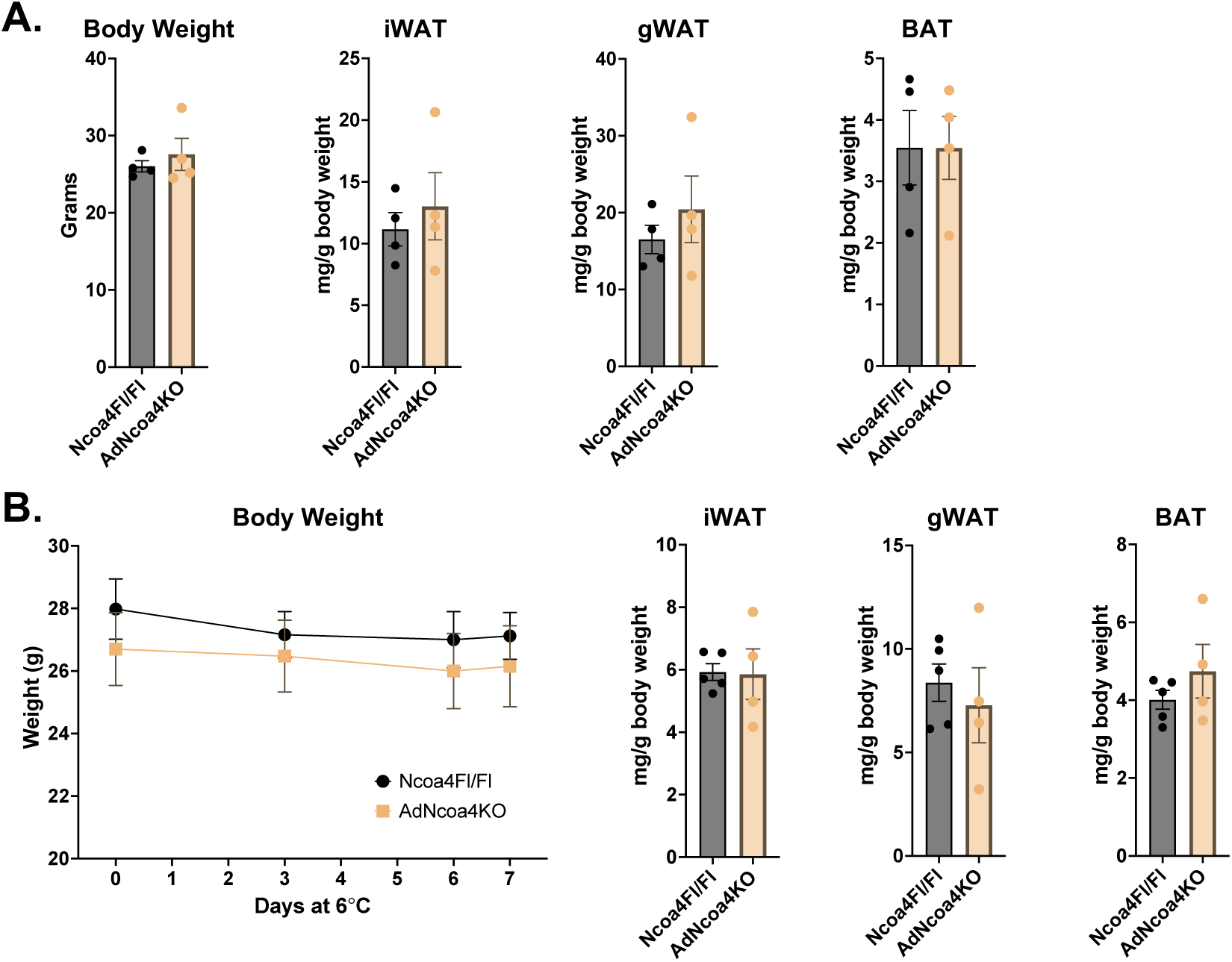
Related to Figure 5. Body weights of AdNcoa4KO mice are similar to controls. A) Body weights and adipose tissue weights 16-week-old chow-fed, room temperature-housed male Ncoa4^Fl/Fl^ and AdNcoa4KO mice. B) Body weights and adipose tissue weights from Ncoa4^Fl/Fl^ and AdNcoa4KO mice housed at 6°C. iWAT = inguinal subcutaneous white adipose tissue (WAT); gWAT = gonadal WAT; BAT = brown adipose tissue.

## References

1. Klein S, Gastaldelli A, Yki-Jarvinen H, et al. Why does obesity cause diabetes? Cell Metab. 2022 Jan 4;34(1):11–20.

2. Czech MP. Mechanisms of insulin resistance related to white, beige, and brown adipocytes. Mol Metab. 2020 Apr;34:27–42.

3. Ahmadian M, Suh JM, Hah N, et al. PPARgamma signaling and metabolism: the good, the bad and the future. Nat Med. 2013 May;19(5):557–66.

4. Lasar D, Rosenwald M, Kiehlmann E, et al. Peroxisome Proliferator Activated Receptor Gamma Controls Mature Brown Adipocyte Inducibility through Glycerol Kinase. Cell Rep. 2018 Jan 16;22(3):760–773.

5. Sugii S, Olson P, Sears DD, et al. PPARgamma activation in adipocytes is sufficient for systemic insulin sensitization. Proc Natl Acad Sci U S A. 2009 Dec 29;106(52):22504–9.

6. Choi JH, Banks AS, Estall JL, et al. Anti-diabetic drugs inhibit obesity-linked phosphorylation of PPARgamma by Cdk5. Nature. 2010 Jul 22;466(7305):451–6.

7. Wang F, Mullican SE, DiSpirito JR, et al. Lipoatrophy and severe metabolic disturbance in mice with fat-specific deletion of PPARgamma. Proc Natl Acad Sci U S A. 2013 Nov 12;110(46):18656–61.

8. Viswakarma N, Jia Y, Bai L, et al. Coactivators in PPAR-Regulated Gene Expression. PPAR Res. 2010;2010.

9. Hauser S, Adelmant G, Sarraf P, et al. Degradation of the peroxisome proliferator-activated receptor gamma is linked to ligand-dependent activation. J Biol Chem. 2000 Jun 16;275(24):18527–33.

10. Qiang L, Wang L, Kon N, et al. Brown remodeling of white adipose tissue by SirT1-dependent deacetylation of Ppargamma. Cell. 2012 Aug 3;150(3):620–32.

11. Yamamuro T, Kawabata T, Fukuhara A, et al. Age-dependent loss of adipose Rubicon promotes metabolic disorders via excess autophagy. Nat Commun. 2020 Aug 18;11(1):4150.

12. Yamamuro T, Nakamura S, Yanagawa K, et al. Loss of RUBCN/rubicon in adipocytes mediates the upregulation of autophagy to promote the fasting response. Autophagy. 2022 Nov;18(11):2686–2696.

13. Sabate-Perez A, Romero M, Sanchez-Fernandez-de-Landa P, et al. Autophagy-mediated NCOR1 degradation is required for brown fat maturation and thermogenesis. Autophagy. 2023 Mar;19(3):904–925.

14. Zhang Y, Goldman S, Baerga R, et al. Adipose-specific deletion of autophagy-related gene 7 (atg7) in mice reveals a role in adipogenesis. Proc Natl Acad Sci U S A. 2009 Nov 24;106(47):19860–5.

15. Singh R, Xiang Y, Wang Y, et al. Autophagy regulates adipose mass and differentiation in mice. J Clin Invest. 2009 Nov;119(11):3329–39.

16. Altshuler-Keylin S, Shinoda K, Hasegawa Y, et al. Beige Adipocyte Maintenance Is Regulated by Autophagy-Induced Mitochondrial Clearance. Cell Metab. 2016 Sep 13;24(3):402–419.

17. Guilherme A, Pedersen DJ, Henchey E, et al. Adipocyte lipid synthesis coupled to neuronal control of thermogenic programming. Mol Metab. 2017 Aug;6(8):781–796.

18. Lodhi IJ, Yin L, Jensen-Urstad AP, et al. Inhibiting adipose tissue lipogenesis reprograms thermogenesis and PPARgamma activation to decrease diet-induced obesity. Cell Metab. 2012 Aug 8;16(2):189–201.

19. Guilherme A, Rowland LA, Wetoska N, et al. Acetyl-CoA carboxylase 1 is a suppressor of the adipocyte thermogenic program. Cell Rep. 2023 May 30;42(5):112488.

20. Henriques F, Bedard AH, Guilherme A, et al. Single-Cell RNA Profiling Reveals Adipocyte to Macrophage Signaling Sufficient to Enhance Thermogenesis. Cell Rep. 2020 Aug 4;32(5):107998.

21. Rowland LA, Guilherme A, Henriques F, et al. De novo lipogenesis fuels adipocyte autophagosome and lysosome membrane dynamics. Nat Commun. 2023 Mar 13;14(1):1362.

22. Han H, Cho JW, Lee S, et al. TRRUST v2: an expanded reference database of human and mouse transcriptional regulatory interactions. Nucleic Acids Res. 2018 Jan 4;46(D1):D380–D386.

23. Fang L, Zhang M, Li Y, et al. PPARgene: A Database of Experimentally Verified and Computationally Predicted PPAR Target Genes. PPAR Res. 2016;2016:6042162.

24. Kajimura S. Promoting brown and beige adipocyte biogenesis through the PRDM16 pathway. Int J Obes Suppl. 2015 Aug;5(Suppl 1):S11–4.

25. Huang J, Linares JF, Duran A, et al. NBR1 is a critical step in the repression of thermogenesis of p62-deficient adipocytes through PPARgamma. Nat Commun. 2021 May 17;12(1):2876.

26. Kollara A, Brown TJ. Expression and function of nuclear receptor co-activator 4: evidence of a potential role independent of co-activator activity. Cell Mol Life Sci. 2012 Dec;69(23):3895–909.

27. Heinlein CA, Ting HJ, Yeh S, et al. Identification of ARA70 as a ligand-enhanced coactivator for the peroxisome proliferator-activated receptor gamma. J Biol Chem. 1999 Jun 4;274(23):16147–52.

28. Mancias JD, Wang X, Gygi SP, et al. Quantitative proteomics identifies NCOA4 as the cargo receptor mediating ferritinophagy. Nature. 2014 May 1;509(7498):105–9.

29. Dowdle WE, Nyfeler B, Nagel J, et al. Selective VPS34 inhibitor blocks autophagy and uncovers a role for NCOA4 in ferritin degradation and iron homeostasis in vivo. Nat Cell Biol. 2014 Nov;16(11):1069–79.

30. Sun W, Dong H, Balaz M, et al. snRNA-seq reveals a subpopulation of adipocytes that regulates thermogenesis. Nature. 2020 Nov;587(7832):98–102.

31. Wang T, Sharma AK, Wu C, et al. Single-nucleus transcriptomics identifies separate classes of UCP1 and futile cycle adipocytes. Cell Metab. 2024 Sep 3;36(9):2130–2145 e7.

32. Bellelli R, Federico G, Matte A, et al. NCOA4 Deficiency Impairs Systemic Iron Homeostasis. Cell Rep. 2016 Jan 26;14(3):411–421.

33. Mancias JD, Pontano Vaites L, Nissim S, et al. Ferritinophagy via NCOA4 is required for erythropoiesis and is regulated by iron dependent HERC2-mediated proteolysis. Elife. 2015 Oct 5;4.

34. Hou W, Xie Y, Song X, et al. Autophagy promotes ferroptosis by degradation of ferritin. Autophagy. 2016 Aug 2;12(8):1425–8.

35. Fujimaki M, Furuya N, Saiki S, et al. Iron Supply via NCOA4-Mediated Ferritin Degradation Maintains Mitochondrial Functions. Mol Cell Biol. 2019 Jul 15;39(14).

36. Suzuki T, Komatsu T, Shibata H, et al. Crucial role of iron in epigenetic rewriting during adipocyte differentiation mediated by JMJD1A and TET2 activity. Nucleic Acids Res. 2023 Jul 7;51(12):6120–6142.

37. Seale P, Bjork B, Yang W, et al. PRDM16 controls a brown fat/skeletal muscle switch. Nature. 2008 Aug 21;454(7207):961–7.

38. Cohen P, Levy JD, Zhang Y, et al. Ablation of PRDM16 and beige adipose causes metabolic dysfunction and a subcutaneous to visceral fat switch. Cell. 2014 Jan 16;156(1-2):304–16.

39. Harms MJ, Ishibashi J, Wang W, et al. Prdm16 is required for the maintenance of brown adipocyte identity and function in adult mice. Cell Metab. 2014 Apr 1;19(4):593–604.

40. Ohno H, Shinoda K, Ohyama K, et al. EHMT1 controls brown adipose cell fate and thermogenesis through the PRDM16 complex. Nature. 2013 Dec 5;504(7478):163–7.

41. Li F, Jing J, Movahed M, et al. Epigenetic interaction between UTX and DNMT1 regulates diet-induced myogenic remodeling in brown fat. Nat Commun. 2021 Nov 25;12(1):6838.

42. Chakravarthy MV, Pan Z, Zhu Y, et al. “New” hepatic fat activates PPARalpha to maintain glucose, lipid, and cholesterol homeostasis. Cell Metab. 2005 May;1(5):309–22.

43. Eguchi J, Wang X, Yu S, et al. Transcriptional control of adipose lipid handling by IRF4. Cell Metab. 2011 Mar 2;13(3):249–59.

44. Yukselen O, Turkyilmaz O, Ozturk AR, et al. DolphinNext: a distributed data processing platform for high throughput genomics. BMC Genomics. 2020 Apr 19;21(1):310.

45. Kucukural A, Yukselen O, Ozata DM, et al. DEBrowser: interactive differential expression analysis and visualization tool for count data. BMC Genomics. 2019 Jan 5;20(1):6.

46. Sherman BT, Hao M, Qiu J, et al. DAVID: a web server for functional enrichment analysis and functional annotation of gene lists (2021 update). Nucleic Acids Res. 2022 Jul 5;50(W1):W216–W221.

47. Huang da W, Sherman BT, Lempicki RA. Systematic and integrative analysis of large gene lists using DAVID bioinformatics resources. Nat Protoc. 2009;4(1):44–57.

48. Zhou Y, Zhou B, Pache L, et al. Metascape provides a biologist-oriented resource for the analysis of systems-level datasets. Nat Commun. 2019 Apr 3;10(1):1523.

